# Mapping the Substrate Recognition Pathway in Cytochrome P450

**DOI:** 10.1101/416305

**Authors:** Navjeet Ahalawat, Jagannath Mondal

## Abstract

Cytochrome P450s are ubiquitous metalloenzymes involved in the metabolism and detoxification of foreign components via catalysis of the hydroxylation reactions of a vast array of organic substrates. However, despite the breadth of cytochrome P450 mediated reactions, a long-standing question is: How does the substrate, in the first place, access the catalytic center of cytochrome P450? The prevalence of conflicting crystallographic evidences of *both* closed and open catalytic center in the substrate-free and substrate-bound cytochrome P450 has given rise to a notion of conformational heterogeneity, which makes the plausible mechanism of substrate recognition by cytochrome P450 puzzling from structural point of view. Here we report multi-microsecond-long unbiased molecular dynamics simulations, which are able to capture the spontaneous process of binding of substrate from bulk solvent to the occluded catalytic center of an archetypal system cytochrome P450cam, at an atomistic precision. In all binding trajectories, the substrate enters through a single channel, where it makes its first contact with the protein-surface and subsequently dwells in a highly long-lived intermediate state, before sliding into the catalytic center of P450cam. The simulated substrate-bound pose and crystallographic pose are in excellent agreement. Contrary to the prevalent hypotheses, our results indicate that a large-scale opening of F/G loop of P450cam is not required for passage of substrate to the catalytic center. Rather, we find that a substrate-induced side-chain displacement of Phe87 residue, coupled with a complex array of dynamical interconversions of multiple metastable substrate conformations along the entry channel, drives the substrate recognition in P450cam. By reconciling multiple precedent investigations, this work put forward an unambiguous view of the substrate recognition mechanism in deep buried cavity of cytochrome P450.

## INTRODUCTION

The cytochrome P450 enzymes are heme-containing monooxygenases, which are one of the largest families of enzymes found throughout the three domains of life with more than 30,000 members sharing a common fold and heme-chemistry.^1-4^ Cytochrome P450s are well known for playing a critical role in both biosynthetic and catabolic processes. These enzymes catalyze a large array of oxidative reactions involved in fatty acid metabolism^5^, steroid hormone biosynthesis^6^, and detoxification of foreign compounds and metabolism of big spectrum of drugs^7^. Because of their ability to catalyze stereo- and regio-selective hydroxylation of hydrocarbons, cytochrome P450s are also potentially attractive catalysts for the degradation of environmental pollutants and for biotechnological drug production.^8^ However, despite the breadth of P450 mediated reactions, a major long-standing puzzle has been: how does the substrate, in the first place, access the catalytic center of cytochrome P450? The current article, using extensive computer simulation, elucidates the comprehensive mechanism of substrate recognition by the catalytic center of the most well characterized P450 system, namely CYP101A1 at an atomistic resolution and identifies dynamic interplay of factors leading the substrate/P450 recognition. Our work maps out the complete pathway of substrate from solvent to the buried catalytic center of cytochrome P450 and unifies multiple prevalent hypotheses relevant to substrate/P450 binding in these contexts.

The CYP101A1^10^, from the *Pseudomonas putida,* known as P450cam, metabolizes camphor (CAM) as a substrate by catalyzing the hydroxylation of D-camphor to 5-exohydroxycamphor. P450cam is the best characterized of all P450s, and in many ways it has served as the archetypal model for all P450s, particularly those with high substrate specificity. However, despite reports of well-resolved crystallographic structures of both substrate-free and substrate-bound form, a dynamic picture of substrate entry from solvent media into the catalytic center of P450cam is still lacking. This is majorly attributed to the fact that the crystal structures of P450cam represent that both substrate-bound^9^, ^11^ and substrate-free^12^ forms are closed conformations, in which substrate-binding site is not readily accessible from the outside. Therefore, it has remained unresolved how the substrate gains access to an active site, which is occluded (see Figure 1A-B). Moreover, the P450cam structures representing the intermediate stages of the catalytic pathways have also been observed in a closed conformation.^13^ The prevalence of these closed conformations in various states of P450cam has raised speculations that during the catalytic cycle, P450cam perhaps opens up to allow the camphor entry, closes it to perform catalysis, and then reopens to allow the release of the product.^14^ The closed conformations of these enzymes during catalysis process are required to control the access of water to the active site to prevent the oxidative damage and the conversion of activated dioxygen to peroxide or superoxide. However, more recent report of first structure of an open conformation of P450cam in the presence of large tethered substrate analogs^15^ and, subsequently, discovery of a crystal structure of open state of P450cam in the absence of CAM^16^, has made the overall picture of substrate recognition in P450cam more complex. The main differences in the open and closed structures were observed in the F- and G-helices and the F/G loop of P450cam. A recent study showed that the structures of P450cam bound to a family of tethered substrates can be divided into at least three distinct kinds of clusters, representing open, intermediate, and closed states^17^ based on the segmental movements of the F- and G-helices. The aforementioned findings, many of which are quite recent, have contributed towards the notion of a “conformational heterogeneity”^18^ in the P450cam. Together, these findings call for revisiting possible mechanism of substrate entry into the catalytic center. In this context, the current article provides an unambiguous atomistic picture into the mechanism of substrate entry into the catalytic center of P450cam, by capturing the complete substrate recognition processes using unbiased molecular dynamics simulations.

**Figure 1:**
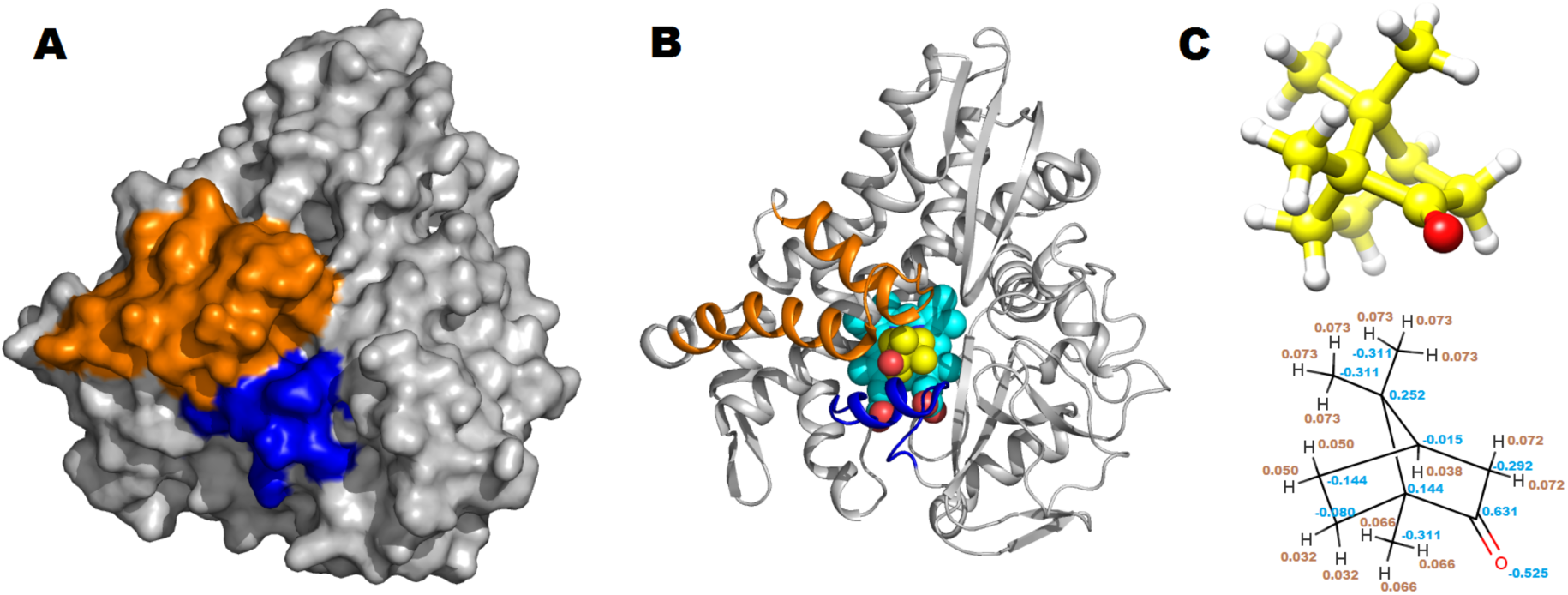
(A-B) Crystal structure of P450cam with camphor bound in heme-containing catalytic center of the enzyme (PDB ID: 2CPP^9^). The space-filling representation indicates that the substrate camphor is bound to a solvent-inaccessible, deeply buried catalytic center. The cartoon representations show the camphor bound to catalytic center. The F, G helices and FG loop region are shown in orange color whereas B’ helix is shown in blue color. The Carbon atoms of heme are shown in cyan color whereas carbon atoms of CAM are shown in yellow color; oxygen atoms of both heme and CAM are shown in red color. (C) Structure of substrate of our interest, camphor. Both licorice and chemical representation is shown. Also embedded in the chemical structure are the partial charges parameterized for this substrate (see model section and SI for detailed set of substrate parameters).

Over the last two decades, multiple attempts have been made to understand the binding mechanism of CAM to P450cam. However, these have been either hypotheses based upon remote readouts of experimental data^9^, ^19-24^ or indirect inferences on substrate-entry mechanisms drawn from *biased* simulation studies^25^, originally designed to simulate the substrate *exit* from P450cam catalytic center. Here, for the first time, to the best of our knowledge, we have been able to capture the spontaneous binding process of camphor entry into the closed and buried catalytic center of P450cam at angstrom level precision, in multiple long unbiased molecular dynamics simulations. The simulated bound pose was found to accurately recreate the crystallographic bound pose. A Markov State Model (MSM) predicts dynamic interconversions of substrate locations along the entry channel, before landing on the catalytic center. These unbiased simulations demonstrate that, contrary to previous hypotheses, complete opening of F/G helix is not required for initial camphor entry early into the pocket. It is discovered that the substrate needs to reside in a long-lived intermediate state (lifetime around 2-3 microsecond), prior to its recognition by the heme catalytic center. We find that a substrate-induced opening of a hydrophobic pocket lined by Phe87 and Phe193 residues drives the overall recognition process. Overall, this work put forward a compelling dynamic picture of the substrate recognition processes in cytochrome P450 and reconciles preceding findings.

## MATERIALS AND METHODS

### Unbiased binding simulation

The X-ray crystal structure of cytochrome P450cam (PDB ID: 2CPP^9^, Resolution: 1.63 Å) without the substrate, was used as the starting point of all simulations. All binding simulations were initiated with the substrate-free form of the P450cam placed at the center of a dodecahedron box with explicit solvent. The minimum distance between the protein surface and the box was 10 Å. The system was solvated with 14618 water molecules and 16 Na^+^ were added to render the system charge neutral. Four CAM molecules were placed at random positions and orientations in the bulk solvent media of the simulation system, which corresponds to a CAM concentration of 13 mM and experimentally suggested 4:1 substrate/protein ratio^26^. It should be noted that there was no direct contact between protein and CAM molecules at the start of the simulation. The final system was having a total 50420 atoms. The all-atom CHARMM36m^27^ force field was used for protein, heme, and ions with the backbone CMAP correction. The CHARMM-compatible parameters for CAM (see Figure 1C) were derived using GAAMP^28^ package (see SI for files with all deduced parameters and coordinates for CAM). All unbiased MD simulations were performed using GROMACS 5.1.4 simulation package^29^, ^30^ using leap frog integrator with a time step of 2 fs. The simulation was performed in NPT ensemble at average temperature of 303 K using the Nose-Hoover thermostat^31^, ^32^ with a relaxation time of 1.0 ps and at 1 bar constant pressure with Parrinello-Rahman barostat^33^ with a coupling constant of 5.0 ps. The Verlet cutoff scheme^34^ was employed with minimum cutoff of 1.2 nm for the Lenard Jones interaction and short-range electrostatic interactions throughout the simulation. Long-range electrostatic interactions were treated by Particle Mesh Ewald^35^ (PME) summation method. All bonds connected to hydrogen atoms were constrained using LINCS algorithm^36^. The bonds and the angles of TIP3P^37^ water molecules were constrained using the SETTLE algorithm^38^. All independent simulations were started from the same configuration by assigning random velocities to all the particles. We carried out three independent multi-microsecond-long unbiased simulations and these simulations were stopped only after the final substrate-bound state was achieved. The substrate-binding process was verified by computing the Root Mean Squared Deviation (RMSD) of CAM between each simulated conformation of MD-derived trajectories and the crystal structure, after removing the rotational and translational motions of protein. The simulation lengths of the binding trajectories were 5.2, 4.9, and 2.7 μs with a total simulation time of 12.8 μs and each simulation was continued well beyond the occurrence of the substrate-binding event. Apart from these long trajectories, many short independent trajectories were initiated from different intermediate states, obtained after clustering of long binding trajectories by considering both protein conformation and substrate location. These short trajectories, with cumulative length of ∼44 μs, were utilized to improve the statistics of underlying Markov state model (discussed later). Overall, an aggregate of about ∼56.8 μs unbiased trajectories were performed. All binding simulations were benefitted by usage of Graphics processing unit (GPU) in the in-house computing facilities.

#### MSM analysis of the MD data

We have employed the PyEMMA^39^ (http://pyemma.org), a Markov state model^40-44^ (MSM) building and analysis package, to identify the kinetically relevant metastable states and their interconversion rates from all the 113 simulated trajectories. The binary contact matrices between protein residues and ligand with the cut-off of 0.5 nm (considering only heavy-atoms) were used as input coordinates. Thus each frame of the trajectories was converted into a row vector with 405 elements, each corresponding to a residue in this 405-residue protein. Further, we employed the time-lagged Independent Component Analysis^45-47^ (tICA) for dimensionality reduction with a lag time of 20 ns. The high dimensional input data were then projected on to the top 12 tICA components based on the kinetic variance for further analysis. Then this 12 dimensional data were used in the k-mean clustering algorithm^48^ to discretize the conformations of all the trajectories into 100 clusters. To find the appropriate lag time, we constructed 100-microstate MSMs at variable lag times. As the implied timescale (relaxation timescale) plot was leveled off at about 5 ns, we considered a lag time of 5 ns to build the final 100 microstate MSM (See Figure S1). A large gap between the second and third eigenvalues in the implied timescale plot suggested that there are three metastable states in the underlying free energy landscape of ligand binding. Thus, a simple three state coarse-grained kinetic model of ligand binding was constructed using hidden Markov model^49^ with a 5 ns lag time. The kinetic parameters (*k*_*on*_ and *k*_*off*)_ were calculated based on the mean-first-passage-time^50^ (MFPT), obtained from the three macrostate MSM. The on-rate and off-rate constants were respectively calculated as *k*_*on*_ = 1/(*MFPT*_*on*_*C*) and *k*_*off*_ = 1*/MFPT*_*off*_ where C is the CAM concentration, 13 mM. Finally, the transition path theory (TPT), as proposed by Vanden-Eijnden and coworkers^51^, ^52^ was used to identify the transition paths and their fluxes. The standard errors were estimated using the bootstrapping method.

#### Free energetics of Phe87-Phe196 gate opening

The free energy surface along the one-dimensional reaction coordinate i.e. distance between the center of mass(COM) of the side chains of residues Phe87 and Phe193 was computed using umbrella sampling simulation^53^. The initial configuration for each window, where CAM is present at various distances between the residues Phe87 and Phe193, was selected from the unbiased binding simulation trajectories, whereas initial configurations without CAM was generated by gradual pulling of the COM distances. The COM distance between the side chains of these residues ranged from 0.45 nm to 1.3 nm with window size 0.02 nm. The windows were restrained by a harmonic potential of force constant 5000 kJ/mol/nm^2^, which allowed sufficient overlap between distributions of adjacent windows. To obtain converged sampling, each of the windows was umbrella-sampled for 20 ns. Then, a weighted histogram analysis method^54^ (WHAM) was employed to obtain the reweighted unbiased potential of mean force along the desired reaction coordinate from the umbrella-sampled windows.

## RESULTS and DISCUSSION

The closed conformation of catalytic center in both substrate-free^12^ and substrate-bound^9^ Cytochrome P450cam with no obvious accessible channel from aqueous solvent, as originally observed in the crystal structure by Paulos and coworkers, prompted us to explore if the spontaneous process of camphor binding to P450cam can be captured by long molecular dynamics simulation. Accordingly, multiple CAM molecules (our substrate of interest) were initially placed randomly in bulk aqueous media around P450cam, by ensuring that there is no direct interaction between the CAM and protein (maintaining a CAM/P450 ratio of 4:1 and CAM concentration of 13 mM)^26^. Subsequently, we initiated multiple independent unbiased all-atom and explicit-solvent MD simulations. By making use of GPU-based computations, each of the simulations was able to access multi-microsecond time scale at reasonable wall-clock time. In three independent simulations of lengths 5.2, 4.9 and 2.7 μs, CAM was able to spontaneously identify the deeply buried heme-binding catalytic center of P450cam (its native binding site) and subsequently achieved correct final binding pose within an angstrom-level resolution. The accompanying movie S1 captures the representative moments defining the CAM recognition by Cytochrome P450 in all three trajectories. All the three unbiased molecular dynamics trajectories show that the distance between binding cavity and CAM gradually decreases as CAM approaches the binding site (Figure 2A, movie S1). The large fluctuations in the distance between ligand and cavity during the initial part of the trajectory, depicts that CAM diffuses in the solvent box extensively and interacts occasionally with the protein, before it identifies the correct entry site on the protein-surface (detailed later). As CAM recognizes the correct entry site, it waits there for a considerably long time at the same place until it slides into the buried cavity to achieve the final bound state. As would be divulged later, the significantly long dwelling of the substrate near the correct entry site, prior to recognizing the native binding site, gives rise to a very crucial intermediate along the CAM recognition pathway.

**Figure 2:**
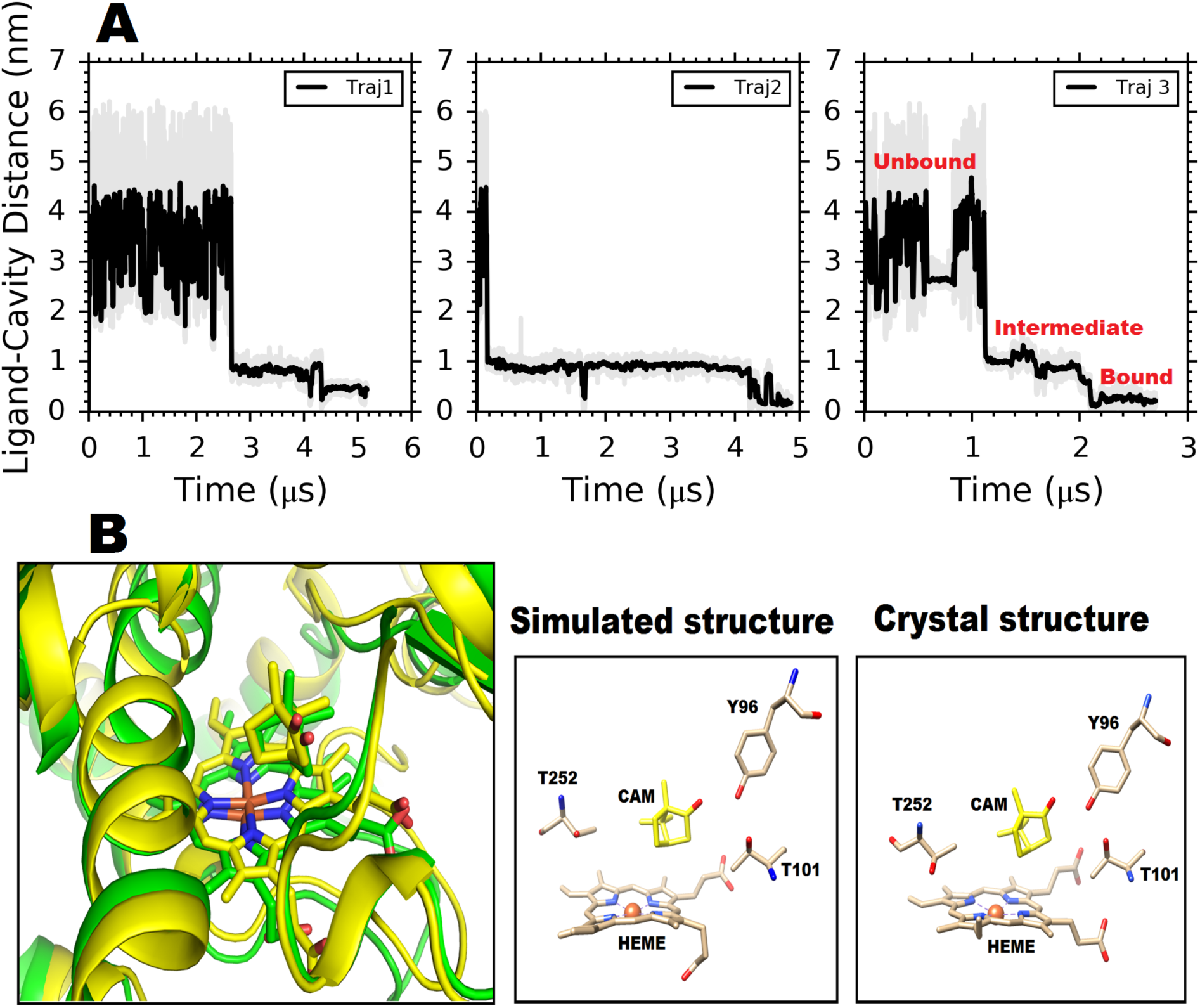
**(A)** Time evolution of distance between P450cam native ‘cavity’ and CAM. Distance profiles from all three successful long trajectories are shown. Protein heavy atoms within 5 Å of CAM in the substrate-bound P450cam are defined as the ‘binding pocket’ or ‘cavity’ (B) Overlay structure of crystal(yellow) and simulation (green) (left side) and comparison of key residue positions inside the binding pocket in both crystal and simulated conformation (right side). The position of CAM and Y96 within hydrogen bonding distance is also prominent in both crystallographic and simulated conformation.

As illustrated by the overlay (Figure 2B) of final substrate-bound structures derived from the MD simulation and crystallography, the difference is almost negligible, suggesting successful simulation of substrate recognition by the P450cam. The close resemblance between crystallographic image and simulated snapshot is striking especially near the catalytic center, with the bound pose of CAM relative to active site residues being very similar in both cases. Both crystallographic image and multiple experimental investigations have earlier reported the existence of a hydrogen bond between Tyr96 and carbonyl oxygen of CAM as a signature of bound pose of CAM.^11^, ^55^ The presence of this Tyr96-CAM hydrogen bond (H-bond) is regarded crucial for regio-selective hydroxylation. As shown by the multiple residue locations near the catalytic center in Figure 2B, our simulated bound structure achieves almost identical crystallographic binding pose inside the pocket by maintaining this specific Tyr96-CAM H-bond, necessary for catalytic activity.

The spatial density profile of CAM around the protein (Figure 3A) (obtained by combining all three long trajectories), presents many locations of proteins, which CAM had explored before entering into a narrow unidirectional channel towards the catalytic center. As depicted by the yellow arrow in the spatial density profile (Figure 3A) and as would be detailed later, this shallow channel would serve as the single link connecting the protein-surface with the catalytic center of P450cam. Nevertheless, the complex nature of MD trajectories associated with CAM binding to P450cam prompted us to build a Markov State Model^41^ (MSM) using our MD trajectories data, to *quantitatively* elucidate the mechanism of CAM binding to active site of cytochrome P450cam in terms of key macroscopic states. Accordingly, we combined the long simulated trajectories, with additional 110 independent short trajectories initiated from the different intermediates from the three long trajectories, (cumulative ∼58 μs trajectories) to obtain statistically meaningful and converged MSM. A simple three-macrostate coarse-grained MSM (Figure 3B) was found to describe the gross kinetic picture of substrate recognition in P450cam quite well (Details provided in method). The MSM-derived *k*_*on*_ and *k*_*off*_ 12 +/- 5 μM^-1^s^-1^ and 203 +/- 68 s^-1^ are found to be in reasonable agreement with experimental values^20^, ^26^ 25 μM^-1^s^-1^ and 21 s^-1^, respectively.

**Figure 3.**
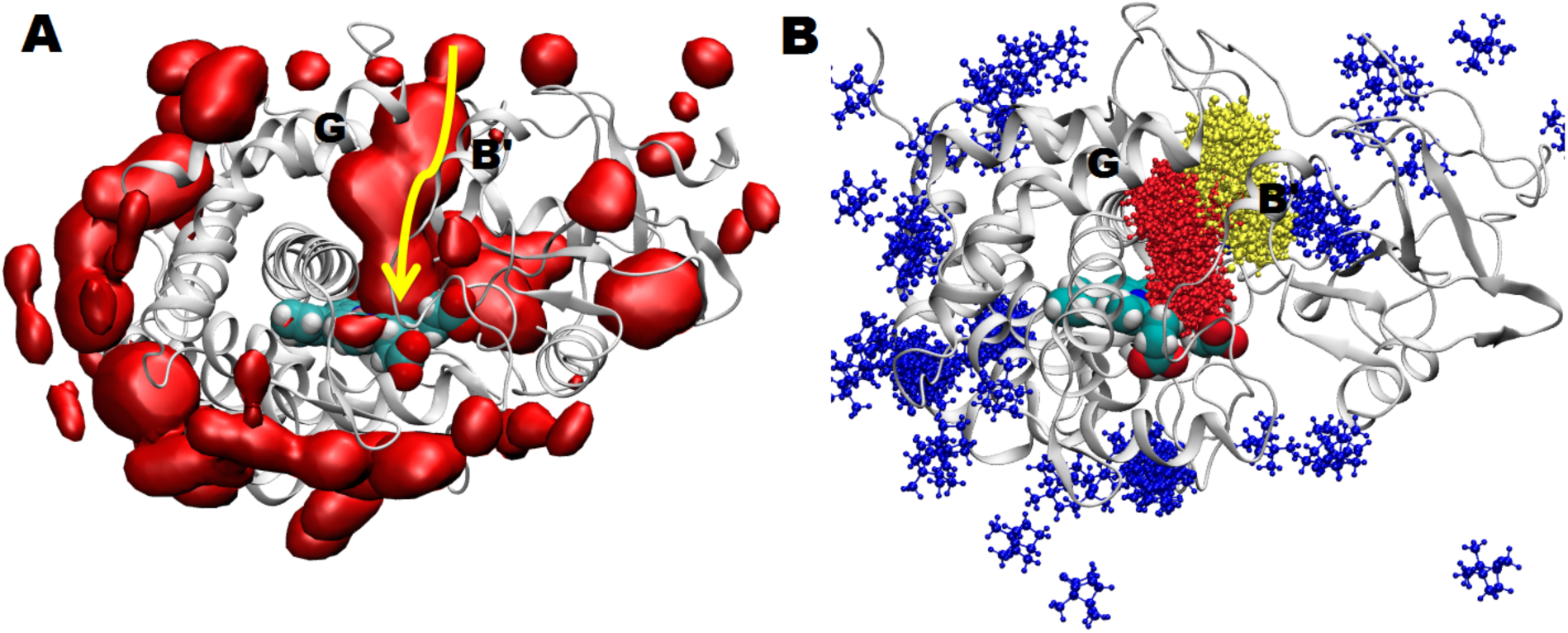
(A.) Spatial density profile of CAM around P450cam. Isosurfaces are shown in red color. Specifically shown, in yellow arrow, is the pathway linking the substrate entry-channel from protein surface to the catalytic center. (B). Ensembles of the three major states of substrate conformations as derived from the Markov State Model. The blue, red and yellow colored macro-states represent the unbound, bound and a long-lived substrate ensemble. 100 snapshots for each of the states are superimposed on the crystal structure of the P450cam, where only CAM molecules are rendered for clarity. Protein is shown in grey cartoon representation and heme is shown in space-filling representations with red for oxygen, cyan for carbon and white color for hydrogen atoms.

Visual inspection of these three MSM-derived metastable states identified these major states as unbound camphor conformation in the aqueous media (blue cluster in Figure 3B), (ii) bound camphor conformation at the catalytic center (red cluster in figure 3B) and (iii) a significantly long-lived intermediate (life time ∼2 μs) at the entry point of F/G helix (yellow cluster in Figure 3B). The analysis clearly indicated *single* dominant pathway for substrate entry via the long-lived intermediate leading to bound pose. An interesting observation from all three long binding simulations was that, the substrate, after exploring all other possibilities, *always* enters into the pocket from the same entry point that is created by the F, G and B’ helix. After visualization of all three trajectories, we observed that, in order to recognize the correct entry site, CAM first contacts with the residues Glu91and Thr192 on the protein-surface. (Figure S2) Quite fittingly, the importance of Thr192 residue in CAM-recognition process has previously been emphasized by mutation-based experiments, where it was found that the rate constant of substrate association is significantly lower in Thr192Glu mutant^56^. Upon making its first contact with the protein surface through Thr192, CAM proceeds along the entry-channel towards the Heme-containing catalytic center. At this stage, the side chains of residues Tyr29, Phe87, Pro89, Pro187, Phe193, and Ile395 and backbone of residues Ala92, Met184, Thr185, Arg186 create a shallow cavity at the surface of protein to hold the incoming substrate. The substrate spends a significantly long time in this small cavity, giving rise to the long-lived intermediate (see yellow ensemble in Figure 3B), (∼2 μs lifetime) as identified by the preceding MSM analysis. The question is: How does the long-lived intermediate, which is unusually stable near the substrate-entry-channel, convert to the native bound state (see red ensemble in Figure 3B) ?

To obtain a close-up view of the mechanism of dynamic conversion of long-lived intermediate to bound conformation along the entry channel, we further rebuilt a higher-resolution MSM than the aforementioned 3-state MSM, this time by selecting *only* the interactions between all the residues lining along this channel and substrate. We repeated the same procedure of MSM building as described in the case of 3-state MSM, except without any further dimensionality reduction and subsequently coarse-grained it into 6-state MSM. Transition path theory was employed to identify the pathways joining the unbound and bound states via different intermediates. The six coarse-grained macrostates are illustrated in Figure 4, where MS1 and MS6 are identified as the unbound and the final bound states, respectively. The macro-states MS2, MS3, MS4, and MS5 are intermediates, each corresponding to CAM being localized at various positions along the substrate-entry channel. As observed by the transition network paths shown by figure 4, these four intermediates MS2-MS5 mutually interconvert in a dynamic fashion, before converting to the native bound conformation (MS6). A closer inspection reveals that the intermediates MS2 and MS3 correspond to macrostate with CAM located near the entry sites at the surface of protein. On the other hand, the intermediate states MS4 and MS5 represent the ensemble of CAM locating inside the binding cavity and are positionally closer to the final binding site. The conformations of MS2 correspond to CAM positioned near the entry site close to the Phe87 and Phe193 opening gate, whereas MS3 is close to the residues Phe26, Met28, and Tyr29. The network of dynamical transitions among the macrostates in Figure 4 also indicates that major net flux from unbound to bound state is initially received by MS2 only. The macrostate MS4, which is slightly inside the pocket and positionally close to MS3, has been observed rarely, when the residues of the entry site create a crevice large enough to allow the dynamic movement of CAM in/out of the cavity. The intermediate state MS5 is very close to the final bound state MS6 and is also directly connected to the Phe87-Phe193 entry gate (detailed later). All the fluxes coming from every intermediate state are finally received by this macro state MS5 before converting into the final bound macro-state (MS6).

**Figure 4:**
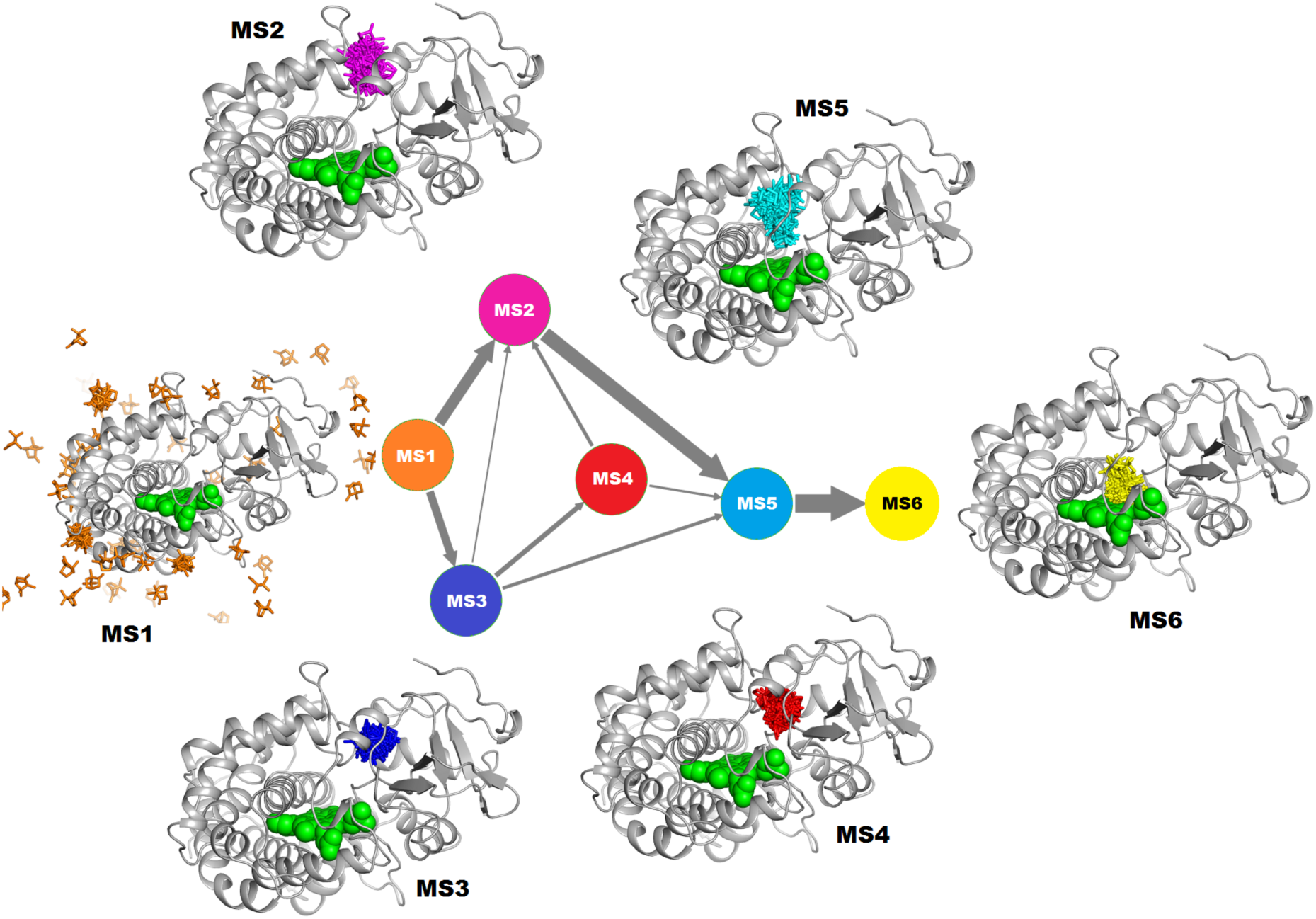
The network kinetically connecting multiple metastable intermediates (MS2, MS3, MS4, MS5) with the unbound (MS1) and bound (MS6) ensemble of CAM along substrate-entry channel of P450Cam. All conformations were derived via building a MSM along the substrate-entry channel of P450cam and the dynamical transitions among these metastable conformations were obtained using Transition Path Theory. Each node of the network represents a macrostate, with the thickness of the arrows denoting the net flux between macrostates. Each coarse-grained macro-state is represented by superimposing 100 snapshots (only CAM molecules are rendered for clarity) and is shown using different colors for CAM. Color scheme for P450cam: White cartoon representation for protein and green space-filling representation for heme catalytic center.

Notwithstanding all aforementioned complex arrays of dynamical transitions among conformations along the substrate-entry channel leading to precise recognition of catalytic center by the substrate, the critical question remains: What specifically drives the substrate-recognition in P450cam? Several cytochrome P450s have been observed to undergo significant conformational changes upon substrate or inhibitor binding. However, it remains unclear whether these conformational changes play an important role in substrate recognition by P450cam. Comparison of substrate-free^16^ (open) and substrate–bound^9^ (closed) crystallographic structures might suggest that P450cam undergoes significant structural change in the region formed by the B’, F and G helices. To understand the role of structural changes in substrate binding process, if any, we calculated the root-mean-squared deviation (RMSD) of Cα atoms of the residues of F, G, F/G loop, and B’ helix, (Figure 5A) from those of the substrate-free closed P450cam structure (PDB id 2CPP). Right axis of Figure 5B shows the time series plot of RMSD of the Cα atoms of residues of region related to large structural deviation in open and close crystal structures (i.e. F, G, F/G loop, and B’ helix, see red colored residue in Figure 5A) and left axis of Figure 5B depicts the time profile of distance between the substrate and the catalytic center for all three trajectories. All trajectories show almost negligible to moderate change in RMSD fluctuations of Cα atoms of the residues of F, G, F/G loop, and B’ helix, which is potentially insufficient for creating large enough crevices for allowing substrate entry into the pocket. In other words, contrary to the hypotheses drawn based upon open and closed crystal structure of P450cam,^9^, ^16^, ^21^ the current work shows that there is no significant opening in F, G and B’ helix region to allow the substrate entry into binding cavity. Previous computational^23^ and experimental studies^24^ had suggested that very strong salt bridge between Lys178-Asp251 and Arg186-Asp251 are crucial for substrate exit/access from/into the binding cavity. These two salt bridges tether the F/G region i.e. F-helix, G-helix, and F/G loop to the I-helix. In our extensively long simulations, we found that the distance between the Cα atoms of Lys186 and Asp251 (Figure S3) does not show large fluctuations, indicating that these salt bridges are not getting perturbed during the binding process. In summary, as reflected in Figure 5, these small-to-moderate RMSD fluctuations of F/G region do not correlate with the changes in substrate-cavity distance profile, especially near the time of substrate recognition and hence do not completely explain the substrate-binding mechanism in P450cam.

**Figure 5.**
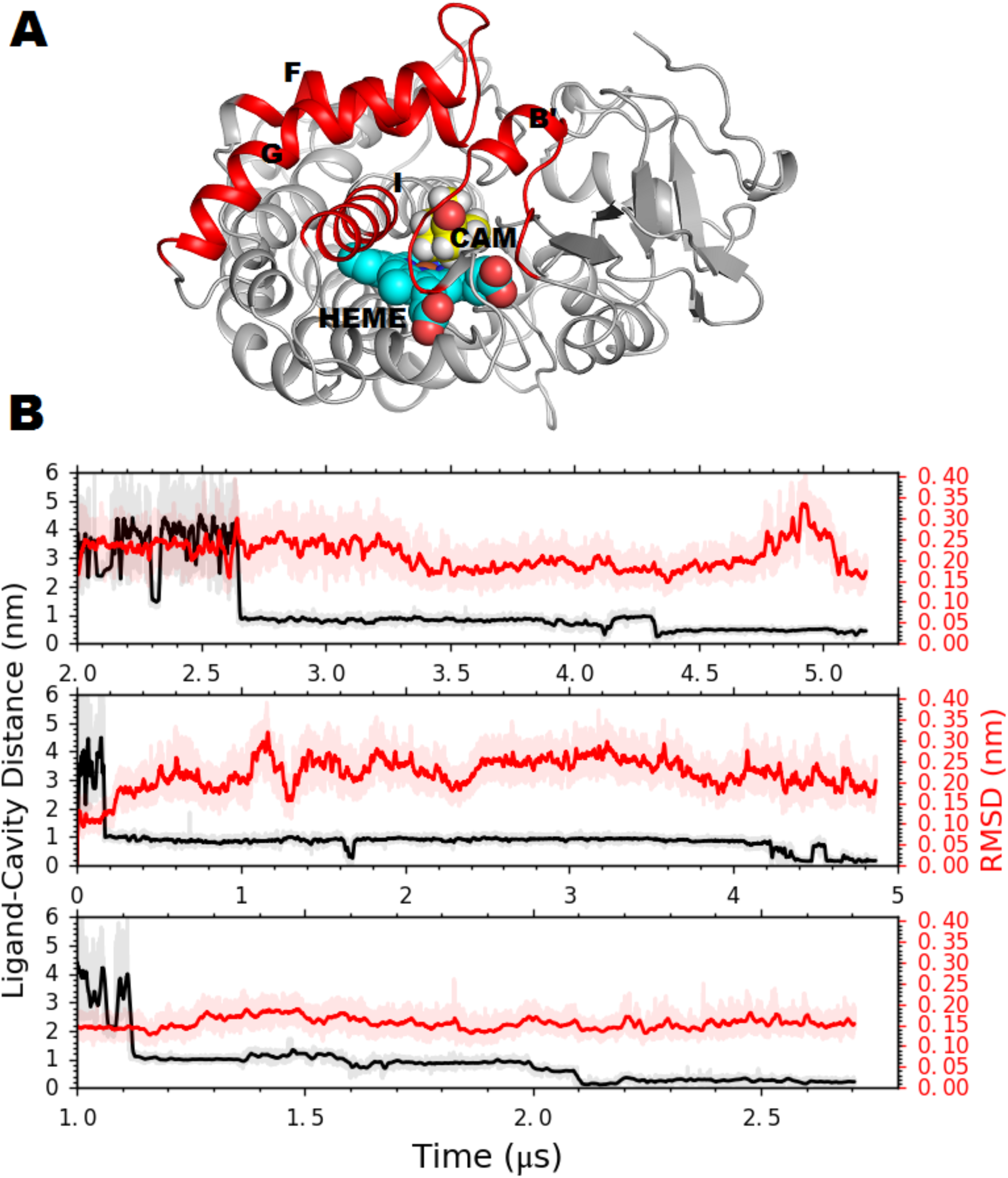
F/G loop opening vs CAM binding. (A) The region of P450cam (F, G, F/G loop, and B’ helix) considered in RMSD calculation are shown in red color, (B) Time series plot of CAM binding to cavity (black color, left axis) and RMSD of residues to measure the F/G loop region opening (red color, right axis).

From the preceding analysis, it becomes clear that CAM binding process does not require significant F/G helix backbone motion for entering into the buried cavity. However, via examination of all MSM-derived macrostates, especially MS2 and MS5, we had observed that the entry site close to the Phe87 and Phe193 opening gate initially holds the substrate from sliding into the native bound conformation (Figure 6A, left). Visual inspection of simulated trajectories reveals that it is eventually the displacement of the side chain of residue Phe87, which had been obstructing the cavity, that creates enough space to allow the CAM to slide into the catalytic center (Figure 6A, middle, right). Accompanying movie S2 demonstrates the process of this crucial event of Phe87-Phe193 gate opening. To demonstrate this, we have calculated the time evolution of distances between Phe87 and Phe193 (Figure 6B). All the three trajectories show that CAM binding is strongly correlated to the distance between Phe87 and Phe193, especially during the time of substrate binding to the catalytic center. As the distance between Phe87 and Phe193 increases to ∼1.0 nm, CAM enters into the pocket to achieve the final bound state (Figure 6A-B). To verify whether the side chain displacement of Phe87 is intrinsic to this protein or actually induced by the binding of substrate, we further employed umbrella-sampling simulations to compute the free energy profile along the Phe87-Phe193 distance (see details in materials and methods). As shown in figure 6C, the reweighted free energy surface along Phe87-Phe193 distance demonstrates that in the absence of CAM, the side chains of these two residues prefer to stay close to each other within 0.6 nm distance and show only one minimum. On the other hand, in the presence of CAM, the location corresponding to free energy minimum gets shifted by 0.1 nm and another second minimum appears at the position of 1.0 nm. The comparison of the energy profiles depicted in figure 6C exhibits that the free energy cost to adopt the conformations that facilitate the entry of CAM into the pocket (i.e.∼1.0 nm distance between Phe87-Phe193, also see middle and right snapshot of Figure 6A) is relatively higher in the absence of CAM than that in the presence of CAM. In other words, this indicates that the probability of the opening of the cavity (i.e. 1.0 nm displacement of the side chain of Phe87) increases ∼28 fold in the presence of CAM, as it is ∼2.0 kcal/mol more unfavorable to increase the distance by 1.0 nm in absence of CAM than that in presence of CAM (e^ΔG/RT^ = 27.7 for ΔG = 2.0 kcal/mol). Therefore, the presence of substrate induces the conformational transition in P450CAM, which implies that the CAM entry into the pocket from entry site might follow the “induced fit” mechanism. These observations are consistent with previous site-directed mutagenesis and stopped-flow kinetic measurement, which had suggested that Phe87_(B/B’ loop)_, and Phe193_(F/G loop)_ play crucial role in substrate access to binding cavity.^20^ Further, the crystal structure of P450cam enzyme with inhibitor containing a long aliphatic tail^21^ had also shown that the orientation of side-chains of residues Phe87_(B/B’ loop)_, Tyr96 and Phe193_(F/G loop)_ was displaced with respect to the camphor bound structure.

**Figure 6.**
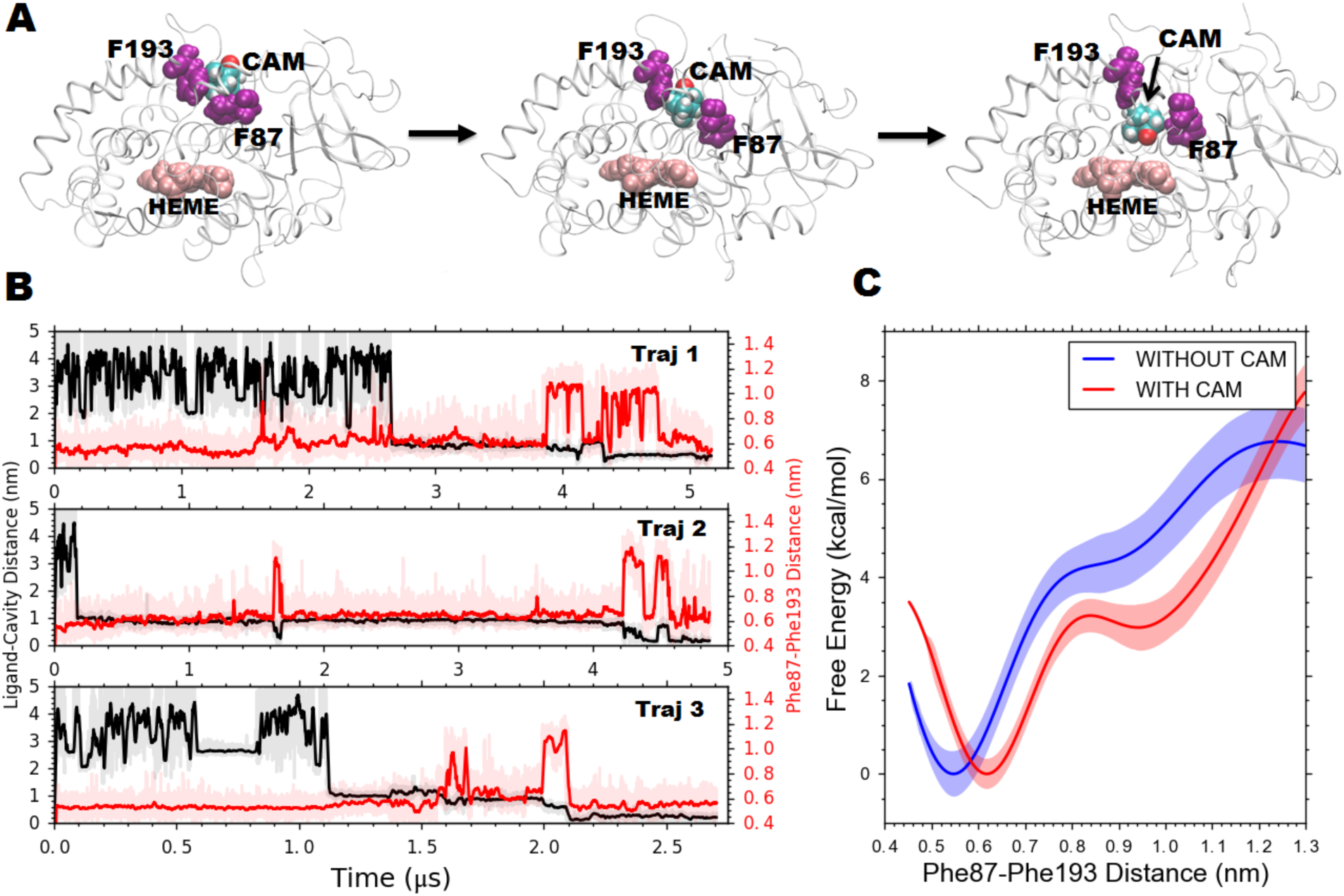
Time series analysis and free energetics of Phe87-Phe193 gate opening. (A) Simulation snapshots to illustrate the different stages involved in Phe87-Phe193 gate opening (also see accompanying movie S2) (B) Comparison of time evolution of CAM binding event and Phe87-Phe193 gate opening, (C) free energy plot along the distance between Phe87 and Phe193 in presence and absence of CAM.

## Summary and Outlook

Despite wealth of experimental and theoretical efforts, the molecular mechanism of substrate recognition by P450s in atomistic detail is not fully understood. Earlier studies of the bound state crystal structure revealed that CAM binds in deeply buried cavity with no obvious accessible channel from aqueous solvent.^9^ Previous theoretical studies based upon biased simulation approach suggested that there are possibilities of multiple entry and exit pathways.^25^ The results of this work, based upon extensive unbiased and unguided MD simulation, put forward a comprehensive mechanism of substrate binding pathway to P450cam. Our multiple unbiased simulations, which spontaneously identify the substrate-binding site at atomistic precision and with overall good binding kinetics, suggest that there is only one substrate-entry path possible, which also has very strong support from previous several experimental site-directed mutagenesis and crystallographic data.

Previous experimental studies have suggested that this protein involves large scale global conformational changes during binding process. However, wade and coauthor’s work^25^ on ligand unbinding pathways showed that the RMSD of Cα atoms of the ‘Pathway class 2’ formed by the residues of F/G loop and B’ helix is overall small and only side chain rotation of key residues is mainly responsible for CAM exit. Our results demonstrated that the substrate binding process does not require the open/close like movements of F/G loop. The binding of substrate at the entry site induces the displacement of Phe87_(B/B’ loop)_ which creates enough space for ligand to enter into the deep buried pocket. Our results are also completely in line with the previous site-directed mutagenesis study^20^ which showed that the substrate binding rate (*k*_*on*_) becomes slower by a factor of 5 if Phe87 is replaced by the larger size residue Tryptophan.

Considering Cytochrome P450’s stature as the key enzyme in the biosynthetic and catabolic processes, the mechanism associated with the crucial substrate recognition stage of P450 catalytic cycle has remained a point of contention. In this context, the atomistically precise spontaneous substrate recognition pathway and related dynamical conformational transitions revealed by the current simulations provides a clear view of the mechanism of the overall process. The insights drawn from this work on conformational flexibility and proposals of induced-fit mechanism for substrate binding keeps the door ajar for future investigations.

## Supporting Information

Graphs illustrating the implied timescales at different lag times for the purpose of building Markov state Model and the time profile of salt bridge fluctuations. Also shown is the representative snapshot of the key residues CAM contacts on P450cam surface. Movies showing the substrate recognition processes in P450cam and the process of CAM-induced Phe87-Phe193 gate opening are enclosed. A zip file containing GROMACS-formatted coordinate and parameters for CAM is also provided.

## Acknowledgments

This work was supported by computing resources obtained from shared facility of TIFR Hyderabad, India. JM would like to acknowledge research intramural research grants obtained from TIFR, India, Ramanujan Fellowship and Early Career Research funds provided by the Department of Science and Technology (DST) of India (ECR/2016/000672).

